# A method for validating the accuracy of NMR protein structures

**DOI:** 10.1101/2020.04.20.048777

**Authors:** Nicholas J. Fowler, Adnan Sljoka, Mike P. Williamson

## Abstract

We present a method, Accuracy of NMR Structures using Random Coil Index and Rigidity (ANSURR), that measures the accuracy of NMR protein structures. It provides a residue-by-residue comparison of two measures of local rigidity: the Random Coil Index [RCI] (a measure of the extent to which backbone chemical shifts adopt random coil values); and local rigidity predicted by mathematical rigidity theory using the computational method Floppy Inclusion and Rigid Substructure Topology [FIRST], calculated from an NMR structural model. We compare RCI and FIRST using a *correlation score* (which assesses the location of secondary structure), and an *RMSD score* (which measures overall rigidity, and mainly assesses hydrogen bond correctness). We test the performance of ANSURR using: (a) structures refined in explicit solvent, which have much better RMSD score than unrefined structures, though similar correlation; (b) decoy structures generated for 89 NMR structures. The experimental NMR structures are usually better, though helical and sheet structures behave differently; (c) conventional predictors of structural accuracy such as number of restraints per residue, restraint violations, energy of structure, RMSD of the ensemble (precision of the calculation), Ramachandran distribution, and clashscore. Comparisons of NMR to crystal structures show that secondary structure is equally accurate in both, but crystal structures tend to be too rigid in loops, whereas NMR structures tend to be too floppy overall.

---

Protein structures are probably the single most important resource for understanding protein function, and are deposited in the Protein Data Bank (PDB), which currently contains around 160,000 structures, of which around 90% are X-ray diffraction structures, 8% are NMR structures, and the rest are mainly from electron microscopy (EM)^1^. The NMR structures are relatively small in number, but are important because they include a high proportion of small proteins with under-represented folds. Most NMR structures are determined in solution, whereas X-ray structures are determined in a crystalline environment. Arguably this makes NMR structures more representative of *in vivo* structures. However, structures are only useful if they are accurate (ie close to the “true” structure) and (equally importantly) can be <u>shown</u> to be accurate. The PDB has therefore become increasingly concerned about validation of structures in the database: the community needs objective and reliable measures to check whether the structure deposited is accurate. The PDB set up four task forces to provide recommendations for validation: for crystallography, NMR, EM and small-angle scattering, which have all reported^2–5^ and have created a suite of validation tools for the PDB^6^. They concluded that validation cannot be based on a single measure. The measures used comprise a combination of geometrical tests, and comparison to input data. Because it is expected that crystal structures and solution structures have the same physical forces underlying them, the *geometrical tests* for crystal and NMR structures are identical, and include clashscore (how well atoms are packed together), an analysis of Ramachandran outliers (how well the backbone dihedral angles comply with structural norms), and an analysis of sidechain outliers. The *comparisons to input data* are necessarily different for X-ray and NMR structures. For X-ray structures there is a very good measure, namely the *R* factor, which is the difference between the intensities of experimental diffraction data, and those calculated from the final structure. If the *R* factor is low (typically less than about 20%) then the structure is almost certainly essentially correct. In structural biology there is a strong temptation to over-fit the data, i.e. to add extra detail in order to improve the fit between experimental data and structure. Hence, a second measure was developed: *R*free, which is an *R* factor calculated using 10% of the diffraction data that was set aside and not used in the refinement^7^. *R*free should be similar in size to *R* for a structure that is not over-refined. Together these two measures provide a reliable guide to the accuracy of the crystal structure.

Unfortunately, no such measure exists for NMR structures^8–11^. The original experimental data have no direct relationship with the structure in the way that diffraction data do, and the experimental input restraints, of which the most common and useful are distance restraints obtained from NOESY spectra, require extensive manipulation and interpretation of the original data before they can be used as restraints. Furthermore, the quantity of information comprising the experimental restraints is far less for NMR. This makes NMR structures inherently less precise, and probably less accurate too, and also means that cross-validation by missing out 10% of the data, as used for *R*free, is not generally possible for NMR structures because we cannot afford to ‘throw away’ 10% of the restraints^12^. NMR structures thus tend to be validated using an unsatisfactory set of restraint comparisons, typically comprising number of restraints per residue, restraint violations, and structure precision (RMS distance between members of the ensemble)^5, 13^. None of these is a direct comparison to the input data, and the third of these is explicitly a measure of precision, not of accuracy, and it is already well established that there is little relationship between precision and accuracy^14–17^.

Hence there is a pressing need to find a better validation measure for NMR structures. Here, we present such a measure. A good validation method should (like the *R* factor) as far as possible compare input data directly to structure. The most obvious input data for NMR structures is the spectra. There have been attempts to do this^18, 19^ but there are major difficulties: there is no good way of accurately calculating chemical shifts from structures; dynamics in solution have big effects on spectra; there are many experimental artifacts in NMR spectra; and the number and variety of input spectra used in structure calculations makes it hard to define or measure what should be compared. Hence, we have here used <u>backbone chemical shifts</u> as our input data. These can usually be obtained reliably and rapidly, and there is little or no manipulation or sorting required, by contrast to distance restraints.

The structure of this paper is that we outline the method before demonstrating how we have validated the method using a range of ‘good’ and ‘bad’ structures and by comparing to other typical measures of structure accuracy. We then demonstrate the power of the method by using it to make comparisons between crystal structures and NMR structures.

## Outline of the method

Backbone chemical shift assignments (ie HN, ^15^N, ^13^Cα, ^13^Cβ, Hα, and C’) can usually be obtained rapidly, semi-automatically, and reliably from a set of triple resonance spectra obtained from^15^N,^13^C double labelled protein. In order to determine a protein NMR structure, shift assignments are the necessary first stage^20^, meaning that any protein that has an NMR structure must have backbone shift assignments (which are now required to be submitted with the structures). Crucially, shift assignments are subject to minimal manipulation. This is very different from distance restraints obtained from NOE spectra. For distance restraints there are inevitably many stages of data sorting and rejection, no matter whether the restraints are inputted manually or automatically. Some person or computer must decide which signals to include, how to assign them, when to reject or modify the restraints, and how to set the calibration between peak intensity and distance restraint. All of these reduce the value of distance restraints as independent quality measures. For all these reasons, backbone assignments are better validation input than distance restraints.

In our method, backbone chemical shift assignments are compared to a structure. Although a number of programs can calculate shifts from structures, they are not sufficiently accurate to perform a useful comparison except in rather general terms^14, 21^. Hence, the heart of our method is that the backbone shifts are used to calculate the *local rigidity* of the backbone, based on an established measure, the Random Coil Index (RCI), which calculates how similar each of the six backbone shifts is to a tabulated “random coil shift” value^22^. It has been shown to provide a remarkably reliable guide to local rigidity, whether measured by NMR relaxation or by crystallographic *B* factor^22, 23^.

We compare local rigidity as predicted by RCI to that computed from a structure using techniques from mathematical rigidity theory. Several software packages and methodologies relying on rigidity theory such as the program Floppy Inclusion and Rigid Substructure Topology (FIRST)^24, 25^ and its various implementations and extensions have been developed for fast computational predictions of rigidity and flexibility of protein structures. Starting with a protein structure, FIRST creates a topological graph (a constrained network consisting of nodes and edges), where atoms are represented by vertices (nodes), and edges represent the constraints corresponding to the intramolecular interactions of a protein e.g. covalent bonds, hydrogen bonds and hydrophobic interactions. Applying the mathematically well-established pebble game algorithm and molecular theorem^26^, FIRST then determines locally rigid subgraphs (rigid regions in the network), a process referred to as *rigid cluster decomposition.* The degree of flexibility can be quantified as a function of hydrogen bond energy by repeating rigid cluster decomposition as edges corresponding to hydrogen bonds are removed incrementally from the graph, and noting the energy at which the Cα atom of a residue no longer belongs to a rigid subgraph, ie becomes flexible. We convert this energy to a Boltzmann population ratio, effectively giving the probability that a residue is flexible.

The two measures of local rigidity (RCI and FIRST) are then compared and a numerical comparison gives a score: a measure of how well the local rigidities match, and thus whether the structures produce a local rigidity that matches the one described by the RCI. Following extensive trials, we use two different measures of similarity: (a) The *correlation* between the two. This tests whether the peaks and troughs are in the same places. Peaks are locally mobile regions while troughs are locally rigid regions, generally regular secondary structure. This comparison therefore mainly shows whether the secondary structure is correct. (b) The *root-mean square deviation* (RMSD) between the two. This tests whether overall the structure is too rigid or too floppy. It is strongly influenced by the geometry of hydrogen bonds and other non-covalent interactions in the structure. As discussed later, the overall rigidity of a structure is determined by not just backbone but also sidechain interactions. Protein structures are often compared by superimposing backbones (often cartoons). Two structures can look very similar in a comparison like this, but one can be much worse than the other in terms of the accuracy of the hydrogen bond network or side chain orientations. In order to assess the relationship between structure and function, it is important that sidechain positions should be correct. The RMSD measure between RCI and FIRST is therefore important because it measures the kind of accuracy needed to interpret function.

Correlation and RMSD highlight different aspects of accuracy, so we decided not to combine them into a single score to represent overall accuracy. Instead, to visualise the overall accuracy of a structure we first calculate the percentile of each measure relative to a reference dataset (as done for various validation metrics published alongside a structure in the PDB), which we term *correlation score* and *RMSD score,* respectively. Then, we plot both on a single graph, as demonstrated in Fig. 1 for four different models of the same protein. The most accurate models (those with good scores for both correlation and RMSD) appear in the top right-hand corner of the plot.

**Fig. 1.**
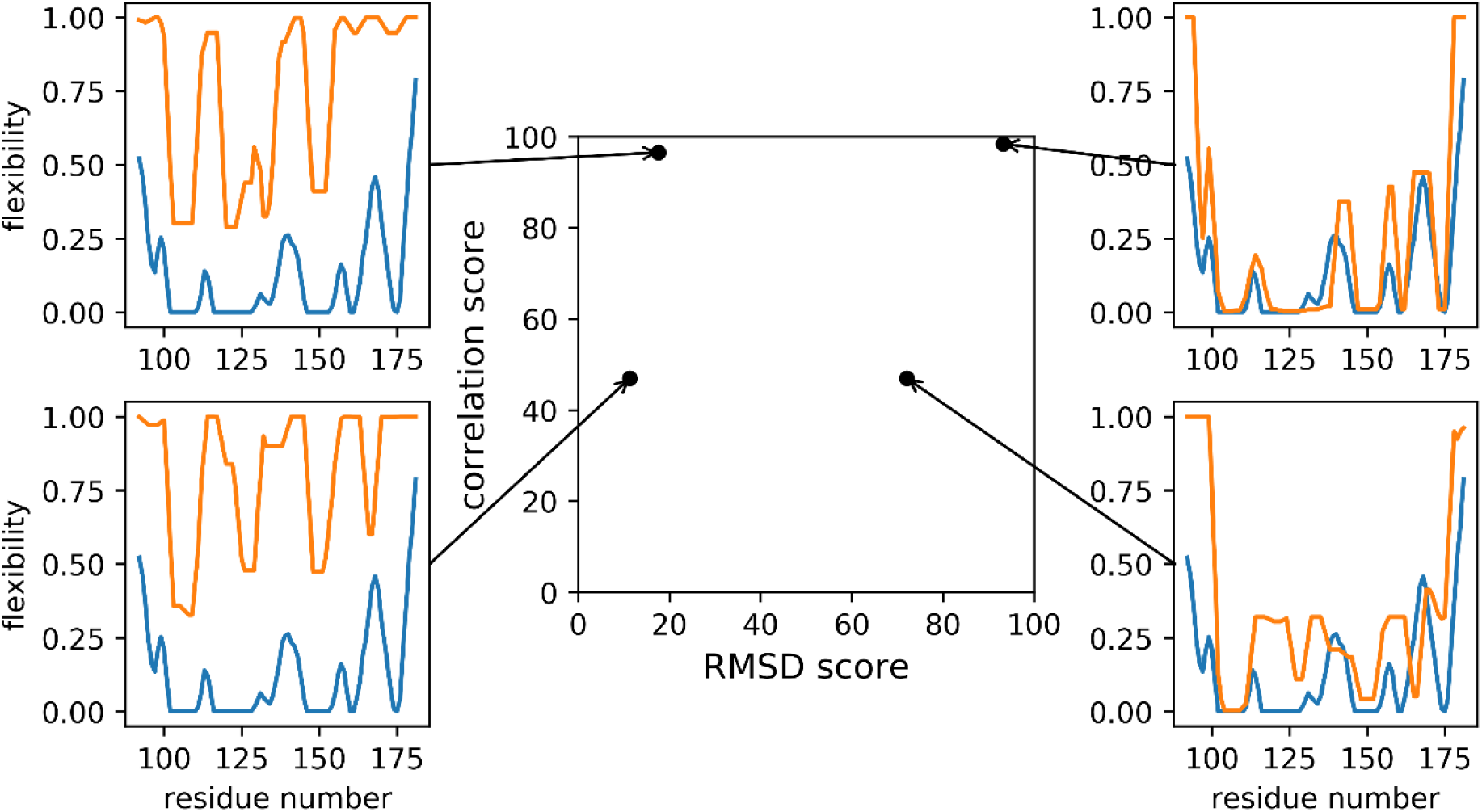
ANSURR analysis of four models from NMR ensembles for the DNA binding domain of the human Forkhead transcription factor AFX (PDB ID 1e17). In the four plots, the blue lines show the flexibility predicted by RCI while the orange lines show flexibility predicted by FIRST. In the center of the figure is the ANSURR analysis showing the RMSD and correlation scores derived from the four models. The two models on the right are from the CNW dataset^27^ (refined in explicit solvent), while the two on the left are from the CNS set (refined *in vacuo).* As is typical, the CNW-refined structures have better RMSD, meaning that the calculated flexibilities compare well on average. The two models at the bottom have poor correlations, because the locations of the peaks do not match well between RCI and FIRST. The two at the top both have good correlations, because the locations of the peaks do match, even though (in the case of the top left structure) their heights are very different.

## Results

There is currently no accepted method for measuring the accuracy of an NMR structure. There are also no databases of “good” or “bad” structures. We have therefore created or adopted datasets that can reasonably be assumed to be bad or good. There are also a range of methods that have been used to measure structure quality, including the geometrical methods described above. We compare our findings to these methods in turn.

### RECOORD CNS (unrefined) vs CNW (refined) structures

The RECOORD project^27^ set out to standardise and tabulate methods for NMR structure calculation. It produced a curated set of structure restraints, which were applied in a consistent manner to more than 500 proteins from the Protein Data Bank (PDB), and then analysed the resultant structures. It carried out two sets of structure calculations on each protein: one using a typical simulated annealing calculation *in vacuo* using CNS (termed CNS) and another using CYANA (termed CYA)^28, 29^. They then took these two sets of structures and refined them in explicit water using ARIA (termed CNW and CYW, respectively)^30^. There is an extensive literature indicating that refinement of NMR structures in explicit water produces better geometries and generally better quality structures^31^, so not surprisingly, the CNW/CYW structures are better.

We have therefore carried out a comparison of those CNS and CNW datasets for which there is sufficient (> 75%) chemical shift completeness, which comprises a set of 173 ensembles each made up of 25 models. From here on we refer to these datasets as CNS75 and CNW75, respectively. In Fig. 2a, the differences in average correlation and RMSD score for each of the 173 ensembles are depicted in a histogram. There is no real improvement in correlation score on refinement in water, with an average improvement of only 1.0. This is expected, as the secondary structure, which ultimately determines the location of peaks and troughs and therefore correlation, changes very little during refinement. As an example, Fig. 2b shows the lack of change in fold for one model. In contrast, RMSD scores are greatly improved, with an average increase of 36.2 and with only one ensemble scoring worse after refinement. This is mostly due to the improvement in hydrogen bonding which acts to rigidify the entire protein. This is easily seen in the difference in computed rigidity before and after refinement (Fig. 2c).

**Fig. 2.**
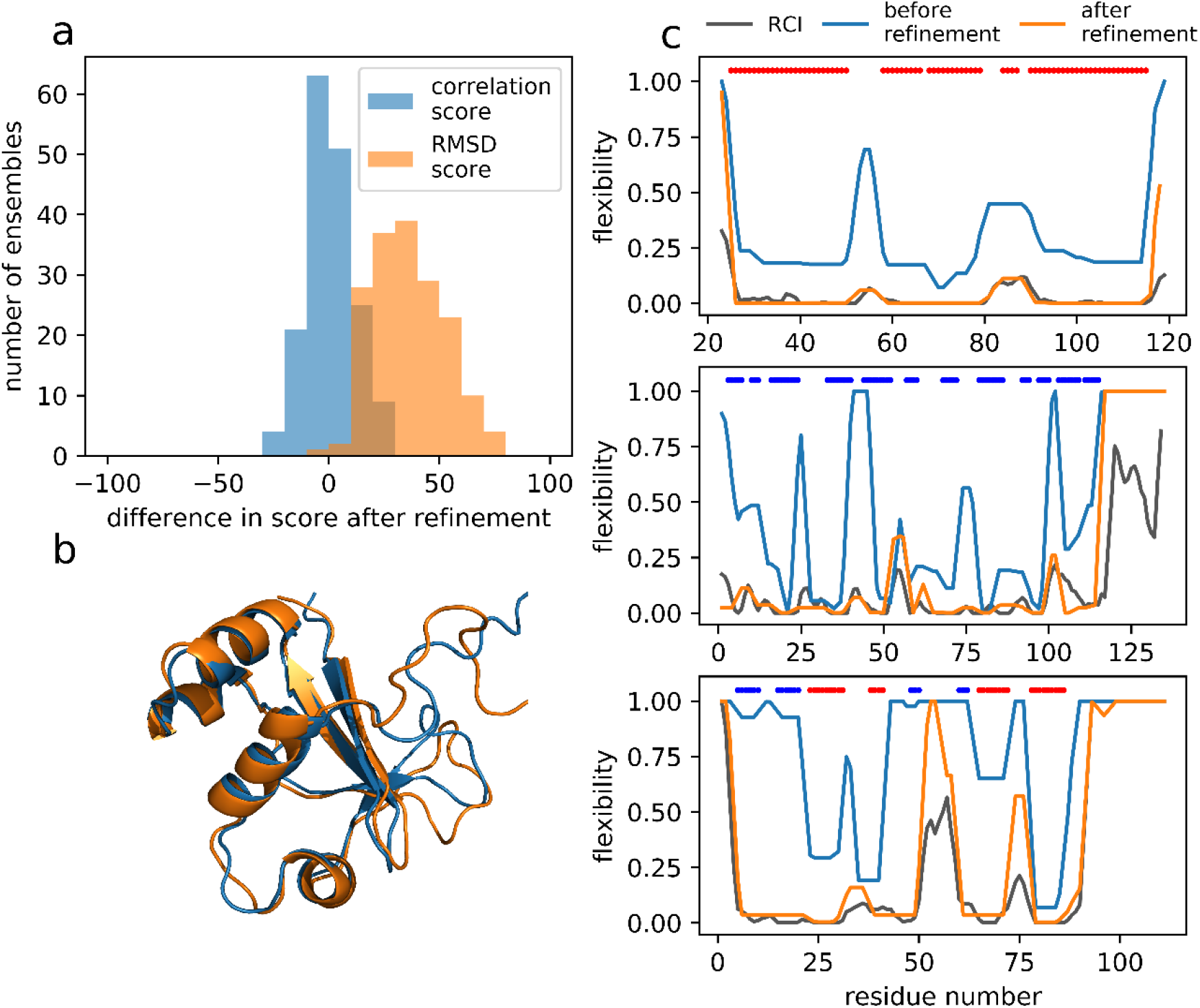
The effect of explicit solvent refinement on the two measures of structure accuracy. (a) Histogram showing the change in average correlation score (blue) and RMSD score (orange), comparing ensembles from the CNS75 to the CNW75 sets. RMSD scores improve dramatically while there is no significant change in correlation scores. (b) Backbone superposition of CNS model 14 and CNW model 14 of the restriction of telomere capping protein 3 from *S. cerevisiae* (PDB ID 1nyn), as a typical example of the effect of refinement in explicit solvent. Although the RMSD score is much better after refinement, the backbones do not look very different. (c) Comparisons of RCI (grey) with flexibility calculated using FIRST for representative models from CNS (blue) and CNW (orange) refinements. The colored bars at the top of each plot show the regular secondary structures: α-helix (red) and β-sheet (blue). The three proteins are (top) the N-terminal domain of VAM3P from *S. cerevisiae* (CNS/CNW model 4, PDB ID 1hs7), a largely helical protein; (middle) a single-domain antibody from *Brucella* (CNS/CNW model 20, PDB ID 1ieh), a largely β-sheet protein, and (bottom) the restriction of telomere capping protein 3 from *S. cerevisiae* (CNS/CNW model 14, PDB ID 1nyn), a mixed α/β protein.

### Decoy vs experimental structures

A straightforward way to generate a pool of structures of varying accuracy is to calculate decoys (physically plausible computationally generated models). We used the 3DRobot web server^32^, which begins from a crystal or NMR structure, identifies possible structure scaffolds from a library, assembles them together, and then refines them, to produce a diverse and well-packed collection of decoys. For about half (79 of 173) of the ensembles in the CNW75 dataset (see Supplementary Information for a list of the chosen models), we calculated a group of 300 decoys. These decoys were then compared to the experimental structure using a Global Distance Test (GDT), which measures the similarity between two structures, calculated as the largest set of Cα in the model structure falling within a defined cut-off of their position in the test structure, after superimposing the structures^33^. A selection of results is shown in Fig. 3a (results for all 79 sets of decoys are depicted in Supplementary Fig. 2). The score for the experimental structure is indicated by a black asterisk and scores for decoys are circles, colored according to their GDT.

**Fig. 3.**
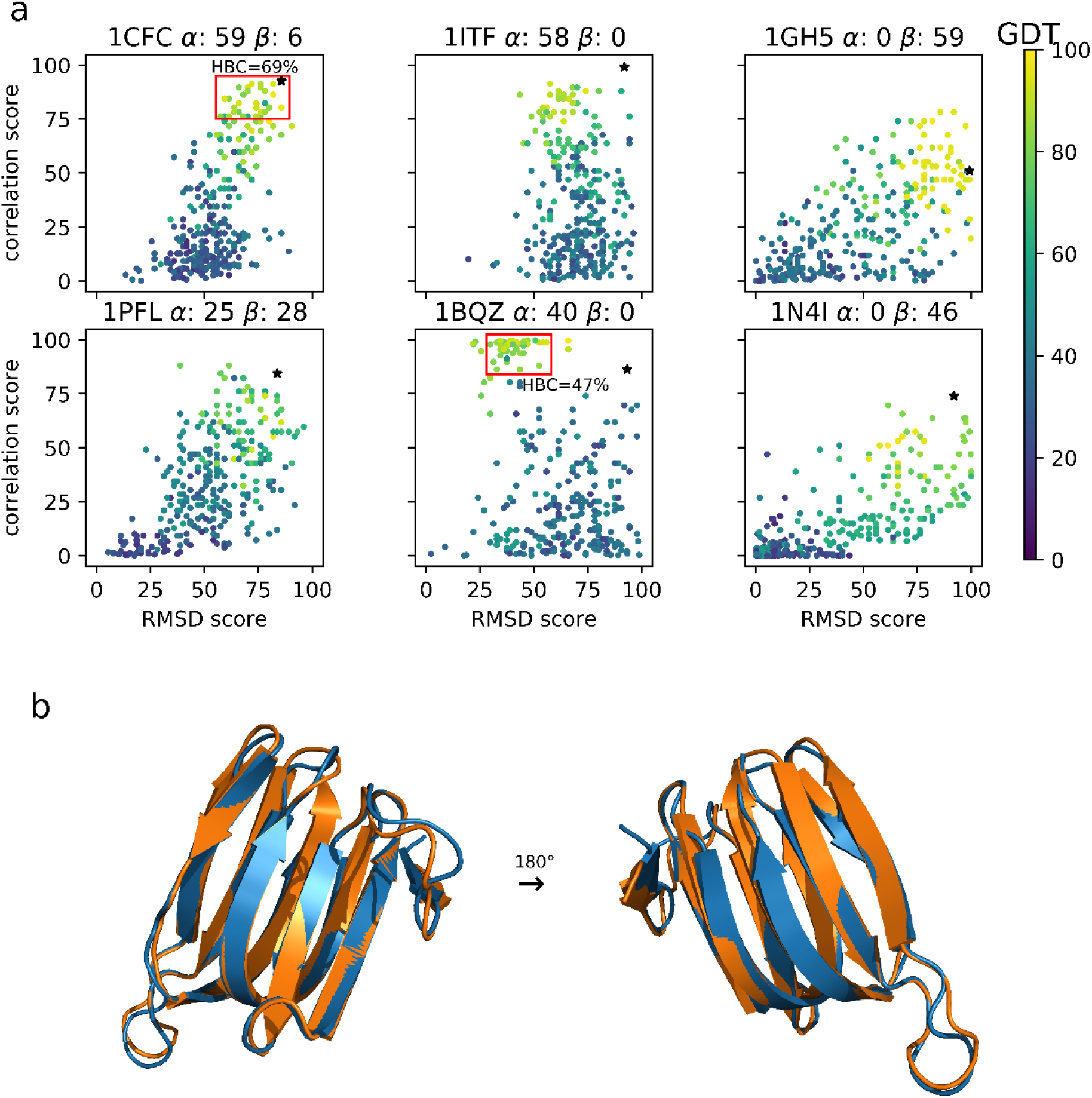
Structural accuracy of decoys. (a) Each plot shows one protein, indicated by its PDB code and the percentage of α-helix and β-sheet in the experimental structure, according to DSSP^36^. The experimental structure is indicated by an asterisk and is the best scoring model in the NMR ensemble, according to our method. The other data show decoys generated by 3DRobot^32^, and color coded by their Global Distance Test (GDT), a measure of similarity to the target^33^, as indicated by the color bar on the right. For two proteins, red boxes indicate the set of decoys used to calculate mean hydrogen bond correctness, as discussed in the text. (b) A comparison of experimental structure (orange) and best decoy (blue) for the protein 1gh5.

From inspection of the examples shown in Fig 3a, it can be seen that the experimental model is usually one of the best structures, as one would expect. Also apparent is that as GDT increases (i.e. as decoys become more like the experimental structure), both the validation scores tend towards those of the experimental structure, confirming that our method does specifically validate accuracy. There is a consistent difference between α-helical proteins (eg 1itf) and β-sheet proteins (eg 1gh5). Helical proteins tend to improve more in their correlation score than in their RMSD score. This seems reasonable: helices are almost always rigid^26^, but not necessarily in the correct location, whereas β-sheet proteins tend to improve more in their RMSD score, because β-sheets can adopt a wide range of local geometries, implying that β-sheet proteins can appear almost correct but have poor hydrogen bonds and thus be much too floppy. Scores for proteins with both α-helical and β-sheet content tend to move in a diagonal, a combination of both effects.

The protein 1bqz presents an interesting example. It is DnaJ, a largely helical protein, and unusually there are many decoys that have a better correlation score but considerably worse RMSD score than the experimental structure, despite most having GDT of around 80 and with some close to 100. However, calculated hydrogen bond correctness scores^34^ i.e. the percentage of hydrogen bonds in the experimental structure that also appear in the decoy, show that these high correlation score decoys (indicated in Fig. 3a with a red box) have poor hydrogen bond geometries (average hydrogen bond correctness of only 47%), and hence a poor RMSD score. By contrast, decoys for 1cfc that approach the accuracy of the experimental structure have good RMSD <u>and</u> correlation score and have better hydrogen bond geometries (average hydrogen bond correctness of 69%).

Another interesting example is the beta-fold protein 1gh5 (an antifungal protein from *S. tendae).* There are some decoys with better correlation and only marginally worse RMSD scores than the experimental structure, suggesting that they are actually more accurate. Fig. 3b compares the experimental structure and best scoring decoy. Immediately obvious (and reassuring) is that at backbone level, both structures are very similar. We note that the experimental structure has a relatively poor correlation score. It is therefore possible that some of the refined decoys genuinely are more accurate: such behaviour has been noted before^35^. Inspection of the full dataset in Supplementary Fig. 2 suggests that this is not uncommon.

### Comparison between ANSURR and conventional NMR-based predictors of accuracy

Conventional predictors of accuracy include the number of restraints per residue used to generate a structure, the number of restraint violations, and the total energy of the structure. The RMSD between models in an ensemble is often used to gauge precision, and by proxy to provide a guide to accuracy. Whilst these measures are expected to be related to accuracy, they do not explicitly determine it. Here we compare these measures to the average RMSD score (Fig. 4a) and correlation score (Fig. 4b) for each ensemble in the CNW75 dataset.

**Fig. 4.**
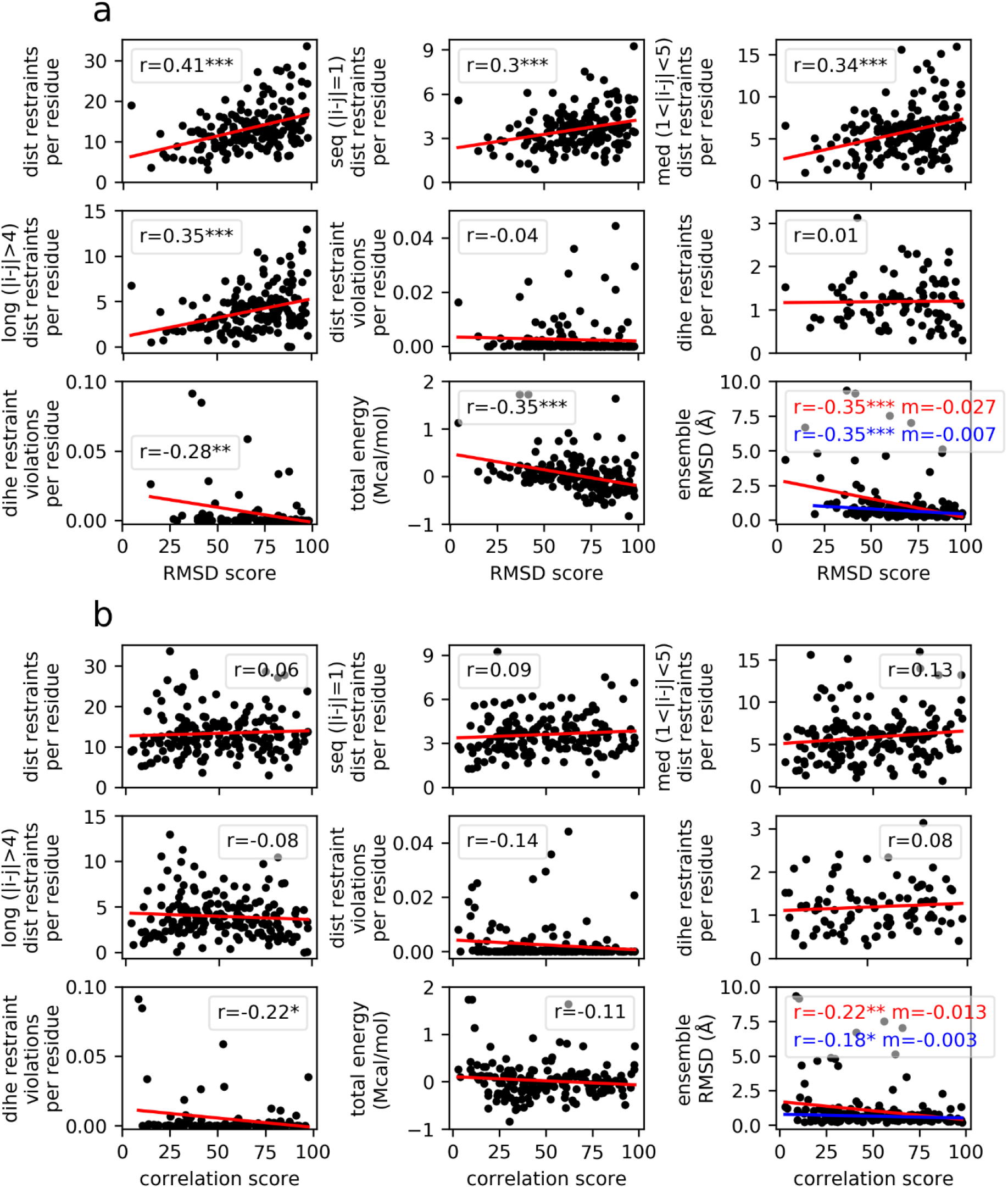
Correlations between conventional NMR-based predictors of accuracy and (a) RMSD score, (b) correlation score. For each plot, the line of best fit and the Pearson correlation coefficient are shown. For the comparisons with ensemble RMSD, fits are shown for all points (red) and for only those points with an ensemble RMSD ≤ 2.5 Å (blue). The statistical significance of the correlation coefficient is indicated by *** *p* < 0.001 ** *p* < 0.01 and * *p* <0.05.

Overall the correlations are much stronger for RMSD score than correlation score. This is not surprising. These predictors largely assess local accuracy, and thus relate to RMSD score better than correlation score.

There is a moderate positive correlation between the number of distance restraints per residue and RMSD score. This is reasonable: a structure with a higher density of distance restraints is expected to be more tightly defined and therefore more (correctly) rigid overall^37^. Categorising distance restraints according whether they are sequential, medium or long-range reveals a slightly better correlation for medium/long-range restraints than for sequential restraints. This is again expected, as medium/long-range restraints provide more information on protein fold, and for this reason are considered a better predictor of accuracy^38^.

The number of distance restraint violations per residue does not correlate with either validation score. Roughly two thirds of structures do not have any violations at all, because structures are normally refined until there are no, or no significant, violations. It is fairly common practice that restraints that are routinely violated during a structure calculation will be discarded along the way. In fact, programs which automate NMR structure calculation do exactly that. For this reason, restraint violations are clearly not a good predictor of accuracy^13, 39 8^.

The number of dihedral restraints per residue does not correlate with either validation score, but dihedral restraint violations do. This is probably because the restraints themselves are relatively weak so that they do not particularly guide the structure to become more accurate. However, weak negative correlation to dihedral restraint violations suggests that these kinds of restraints successfully flag major issues.

There is a moderate negative correlation to the total energy of the structure. Typically, the selection of the final set of structures to represent the ensemble is based on total energy, and the correlation seen here suggests that this is a reasonable way of identifying good structures.

Both RMSD score and correlation score are negatively correlated with ensemble RMSD suggesting that more precise ensembles do also tend to be more accurate. However, if those ensembles with RMSD larger than 2.5 Å are excluded (blue fit lines) then the gradient becomes almost zero, suggesting that for better structures, ensemble RMSD is a poor guide to accuracy. Similar comments have been made previously^14–17, 40^.

In summary, our measures of accuracy match reasonably well to expectations: the number of distance restraints per residue is a fairly good predictor of accuracy, while dihedral restraints, and distance and angle violations, are not. Precision (ensemble RMSD) is a poor predictor of accuracy, while overall energy is surprisingly good as a predictor of accuracy.

### Comparison between ANSURR and geometry-based validation measures

It is unclear whether a correlation should be expected between geometrical quality and accuracy. However, given that NMR structure calculation is to a large extent an optimisation of models, using both NMR-derived restraints and knowledge-derived geometrical factors simultaneously, it is reasonable to expect that an accurate structure should also have good geometrical quality. We therefore compared our validation scores with two widely used indicators of geometrical quality: Ramachandran outliers and clashscore^41^. The program ramalyze (part of the Molprobity suite of validation tools) was used to compute the φ/ψ angles for each residue in the CNW75 dataset and categorise them as either favorable, allowed or outlier. The program clashscore (also part of Molprobity) was used to compute the average number of clashes per 1000 atoms for each ensemble in the CNW75 dataset. In Fig. 5a and 5b, the results for each ensemble are plotted against RMSD score and correlation score, respectively.

**Fig. 5.**
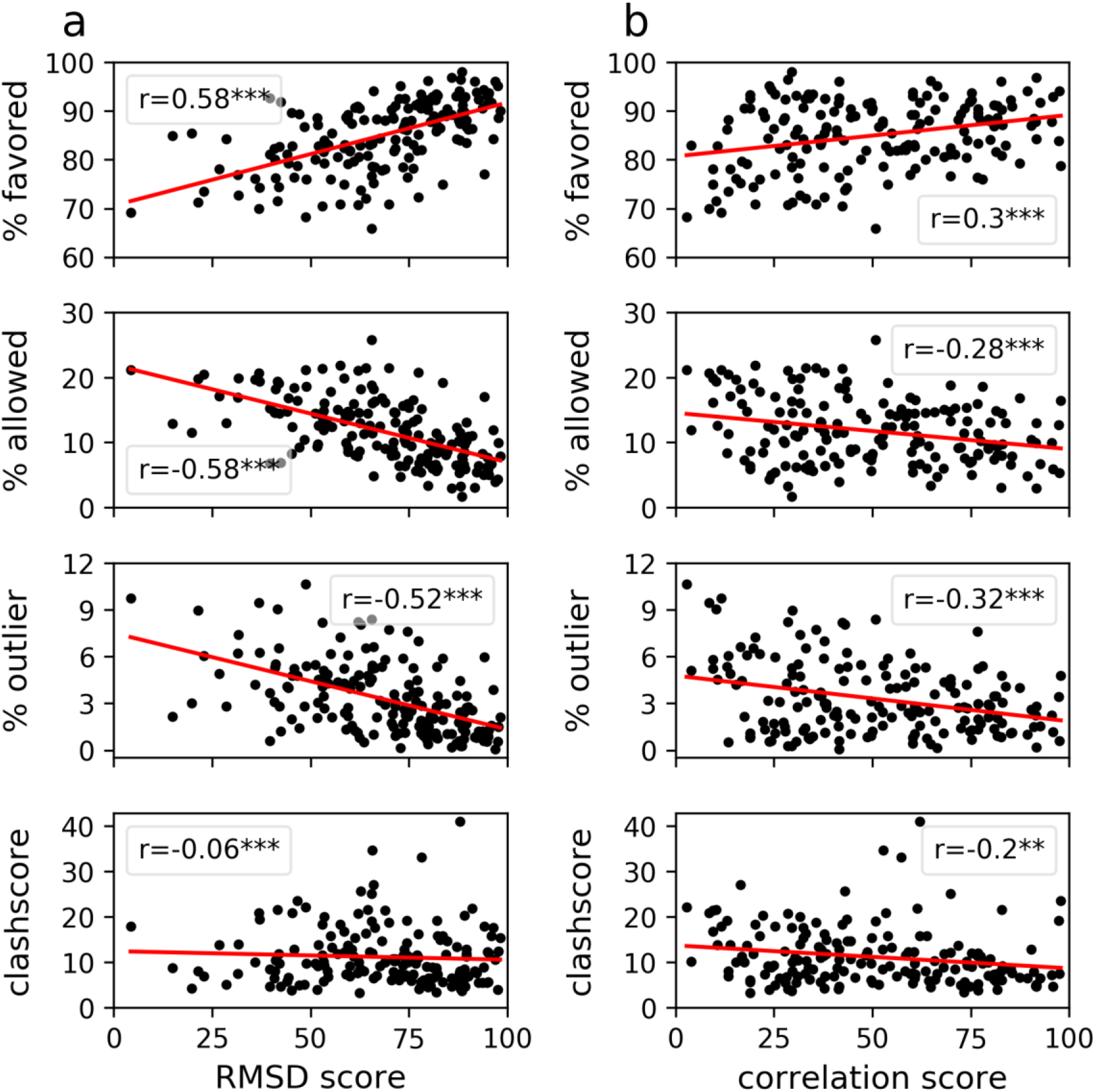
Correlations between geometry-based measures and (a) RMSD score, (b) correlation score. The statistical significance of the correlation coefficient is indicated by *** *p* < 0.001 ** *p* < 0.01 and * *p* <0.05.

The correlation between Ramachandran distribution and RMSD score is the best for any of the measures presented here. In other words, an ensemble with good Ramachandran distribution (high percentage in the favored category, low percentage in the additionally allowed category, small percentage in the outlier category) is likely to have good accuracy. It seems reasonable to find that the most accurate structures are in general those with the best backbone geometry, as was proposed many years ago^42^. By contrast, the comparison with clashscore is weaker. Simulated annealing structure calculations typically use van der Waals repulsions but no van der Waals attractions in the force field: they also have explicit hydrogen atoms in the calculations. It is therefore possible that the NMR-derived restraints are not sufficiently strong or numerous to compete well against van der Waals repulsions, which are therefore effective ways of minimising clashes.

### Comparison of NMR and x-ray crystal structures

An obvious first test for this method is to compare NMR and X-ray crystal structures. It is important to stress here that because we compare the structures to time-averaged chemical shifts obtained using solution NMR, we are explicitly testing how well the structures compare to the average state of the protein *in solution.* Crystal structures are almost always based on many more experimental values, and more precisely measured values, than NMR structures. One would therefore inherently expect them to be more accurate, except that crystal structures represent the structure of the protein in a crystalline environment, whereas the NMR chemical shifts measure structural rigidity in solution. We are therefore here making a somewhat unfair, but important, comparison, namely how well X-ray structures represent the structure of a protein *in solution.*

Here we compare X-ray structures for 68 proteins taken from the set used to train the SHIFTX2 program for predicting chemical shifts^43^ with corresponding NMR structures taken from the PDB (see methods for details). We validated each structure using our method and averaged the validation scores over each chain for X-ray structures, and each model for NMR ensembles. The results are shown in Fig. 6. The correlation scores for X-ray and NMR structures are very similar. In other words, the locations of rigid and flexible regions, generally representing regular secondary structure in solution, are calculated similarly well by both methods. The slightly lower correlation score for X-ray structures originates from some loops seeming to be too rigid. That is, X-ray structures are missing some peaks in flexibility that should be there according to RCI. Crystal structures are obtained from crystalline arrays, and are usually obtained at cryo-temperatures, both of which will tend to reduce the observed flexibility. There is a large body of evidence^44–46^ that crystal structures obtained at room temperature show much more local variability than do structures obtained at cryo-temperatures, and calculations on lysozyme confirm that the room temperature structures have flexibility that matches the RCI data much better than cryo-temperature structures (Supplementary Fig. 3). By contrast, in the RMSD score comparison, on average crystal structures are significantly better. When one inspects the data for individual proteins, it is clear that NMR structures are in general much too flexible, particularly in loop regions. This is not unexpected, as NMR structures often have few restraints in loops.

**Fig. 6.**
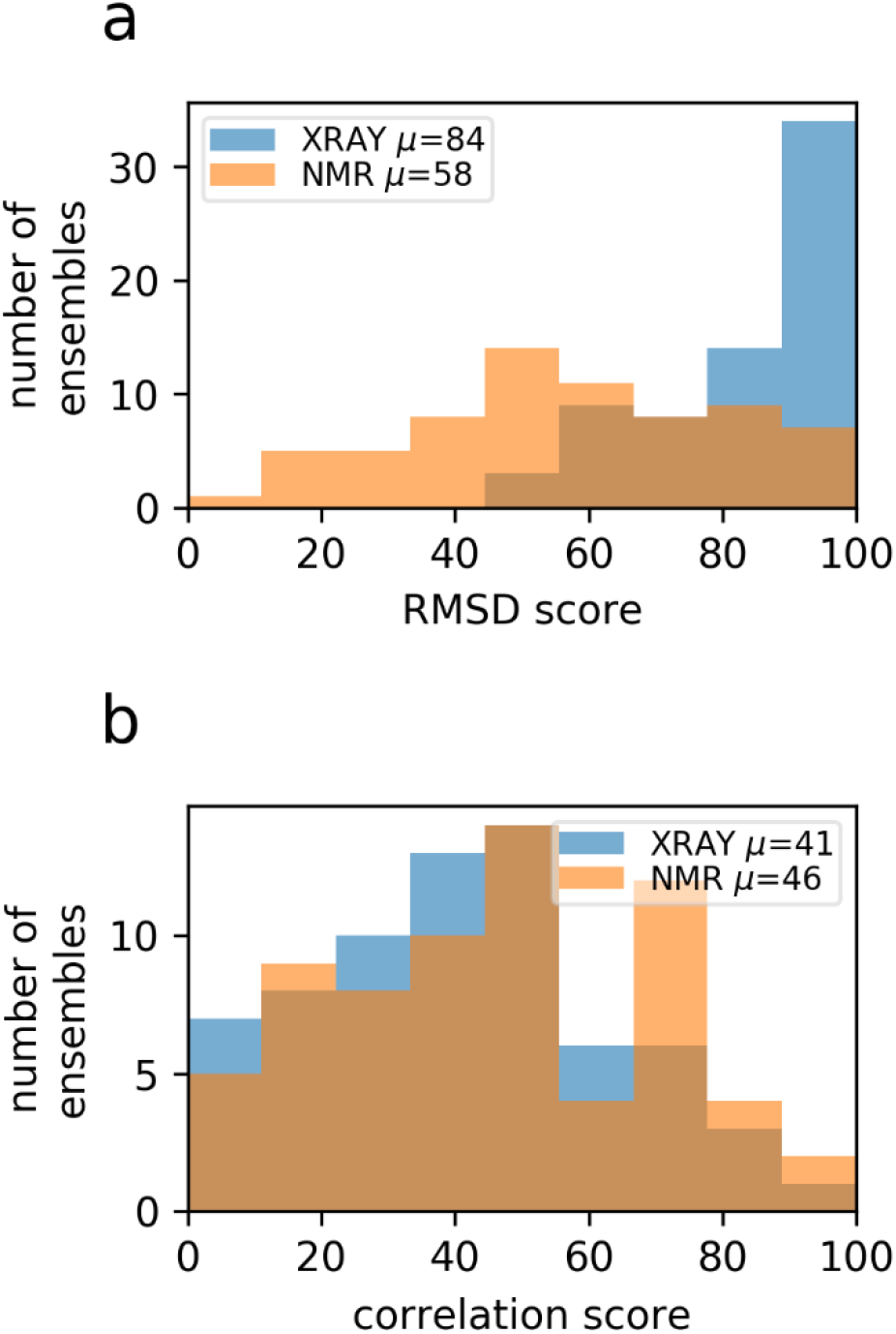
Frequency distributions for X-ray structures (blue) and NMR ensembles (orange) as a function of (a) RMSD score and (b) correlation score. Overlapped distributions are shown in brown. The mean values for each score are shown in the inset box.

## Discussion

We present a new method for determining the accuracy of NMR structures. A range of methods have been proposed previously^10, 13, 42, 47^, including various attempts at an NMR *R* factor^18, 19, 48–50^. Our method has the merits of being simple, rapid, and in agreement with intuitive expectations. Considering that the first NMR structure of a globular protein was published in 1985^51^ it is remarkable that it has taken this long to come up with a workable measure. The lack of a good measure of accuracy has inhibited researchers from using NMR structures; it is hoped that this method will give users more confidence in the use of structural data from NMR.

Because there are no general methods for measuring accuracy, and thus no agreed sets of “good” or “bad” NMR structures, we have been forced to create our own comparisons. Similarly, there are a range of measures that have been proposed for measuring accuracy. In particular, the PDB NMR validation task force^5^ has recommended a set of measures, combining geometrical comparisons and comparisons to input data. These measures are investigated here. We find that the best indicator of accuracy is a Ramachandran analysis, using either the proportion of residues in the favored region or the proportion of outliers. We find that the RMSD between models in an ensemble is a poor measure of accuracy (though an excellent measure of precision, reinforcing the concept that accuracy and precision are largely independent). Other common restraint-based measures of accuracy, such as restraints per residue^8^ or restraint violations, are also poor measures of accuracy^52^. We suspect that part of the problem is that the route from NOE spectrum to distance restraint contains a large number of user-defined decisions (many of which are increasingly being made by the programs, and are thus becoming even more opaque), so that the link between spectrum and restraint is ill defined.

An interesting conclusion to come from this comparison is that the most common measure of structural similarity, backbone RMSD, misses many of the interesting differences. Structures can look very similar when superimposed on the backbone, but contain large variability in sidechain position and hydrogen bond geometry, which has major impact on docking algorithms and on functional aspects such as allostery, enzyme catalysis^53^ and dynamics.

Now that we have a reliable measure of accuracy, it can be applied to some key problems, for example: (1) how good are the NMR ensembles in the PDB? (2) Can we determine which structures in an ensemble are good, and which are not, and can we therefore improve the ensemble? (3) Is it possible to use experimental NMR data to validate or refine protein structure prediction methods? (4) Can one use these methods to identify local errors in NMR structures? We plan to address these questions in the future.

## Methods

### Random Coil Index (RCI)

RCI quantifies local (i.e. per residue) protein flexibility by calculating an inverse weighted average of backbone secondary chemical shifts. We calculate RCI essentially as done by Berjanskii and Wishart^22^, though with a few differences. In the originally published method, the weighting coefficients were not normalised. That is, the sum of the weights for different combinations of shifts did not add up to the same value and therefore the baseline rigidity measure could vary when comparing RCI values calculated with different combinations of shifts. We addressed this by simply dividing the sum of weighted secondary shifts by the sum of the weighting coefficients. We therefore compute RCI as:

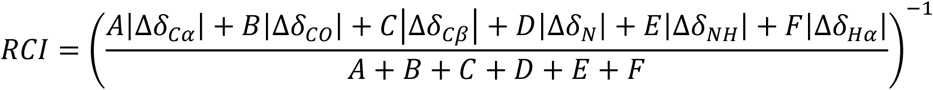

where the Δδ_*l*_ are secondary chemical shifts and *A-F* are weighting coefficients. Some nuclei (Cα, Cβ) are more descriptive than others (HN, NH) and so have larger weighting coefficients. Missing chemical shifts have a weighting coefficient of zero. Another difference is that we use random coil values and nearest neighbor sequence corrections using data obtained from intrinsically disordered proteins^54^ rather than data based on unfolded peptides or proteins (eg^55^). A result of these differences is that our approach outputs a value between 0 – 0.2, rather than between 0 – 0.6 as in the originally published method.

We use the set of optimised weighting coefficients for each of the 63 different combinations of backbone chemical shifts as found in the downloadable Python version of RCI http://www.randomcoilindex.com/. For some combinations, we found the similarity between flexibility predicted by RCI and FIRST is significantly decreased suggesting that, in these instances, RCI is a poor predictor of flexibility. Ultimately, the most reliable validation scores are obtained when a full complement of backbone chemical shifts are provided. Our method will allow validation with any combination/completeness of shifts but the resulting validation score is flagged as less reliable if total chemical shift completeness drops below 75%. For proteins with sufficient chemical shift completeness (>75%), we assume that residues with completely missing backbone chemical shift assignments are missing because the residues are highly mobile. We assign such residues a secondary chemical shift of zero (i.e. they are assumed to be entirely random coil-like) prior to 3-residue smoothing. However, these data points are not used when calculating validation scores.

### Floppy Inclusion and Rigid Substructure Topography (FIRST)

Given a protein structure, FIRST^25^ generates a graph (constraint network) composed of vertices (nodes), which represent atoms; and edges, which represent constraints imposed by the local geometry. Single covalent bonds are modelled by five edges between bonded atoms; double bonds by six; hydrophobic interactions, which are less geometrically constraining, by two; and hydrogen bonds by between one and five, depending on how one chooses to model them. Overall this multigraph represents a generic realization of a molecular body-bar framework in rigidity theory^26^. Typically, rigidity analysis is performed at a range of hydrogen bond energy cut-off values, where hydrogen bonds that meet the cut-off threshold are assigned five edges wile weaker interactions are ignored.

Atoms are considered to be rigid bodies each with six degrees of freedom (three position and three orientation). These degrees of freedom are removed as constraints are added between them. One edge removes up to one degree of freedom e.g. a single covalent bond can remove up to five degrees of freedom between the two bonded atoms. FIRST then uses the combinatorial pebble game algorithm (which checks the counting condition prescribed by rigidity theory^56^) to rapidly decompose the graph into maximum rigid clusters and flexible regions, a process known as rigid cluster decomposition. We consider a residue to be rigid if the Cα atom belongs to a rigid cluster that contains at least 30 atoms: this is a useful caveat because it prevents prolines and aromatic residues automatically showing up as rigid.

Relative flexibility is quantified using a process termed hydrogen bond dilution, which is analogous to the thermal denaturation of a protein. Dilution involves incrementally removing edges associated with hydrogen bonds in the graph (weakest to strongest), repeating rigid cluster decomposition and noting the hydrogen bond energy at which the Cα atom of each residue is no longer part of a rigid cluster i.e. becomes flexible. We have adapted this slightly, choosing to convert the energies to a Boltzmann population ratio at 298.15 K to represent the probability that a residue is flexible.

### Comparing RCI and FIRST

A simple comparison of RCI and FIRST is not ideal, because the frequency distributions of RCI and FIRST output values are different (Supplementary Fig. 1a,b). The main difference is that RCI is calculated as the inverse of averaged secondary chemical shifts and therefore it is not possible to achieve a RCI value of zero. We decided to rescale RCI values so that the mode RCI value (0.024) becomes “zero” and round up any subsequent negative values. At the other end of the scale, particularly noticeable is a large spike in RCI values at 0.2 which is comprised of terminal residues. A similar spike, also comprised of terminal residues, is present in the frequency distribution of FIRST at Boltzmann population ratio equal to one (i.e. completely flexible at 298.15 K). We therefore decided to scale RCI values so that these spikes align. Subsequent values above one (i.e. apparently more flexible than terminal residues) are rounded down, although such instances are very rare. The equation below outlines how we compute rescaled RCI (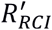) from the original RCI values (*R_RCI_*):

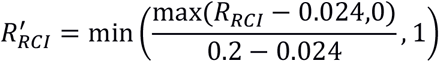

Comparing the frequency distribution of the rescaled RCI and FIRST output values shows good agreement (Supplementary Fig. 1c,d).

### Validation scores

RCI and FIRST are compared using two different measures. One is the correlation, calculated using a Spearman rank correlation coefficient. The other is the root mean square deviation (RMSD), calculated as:

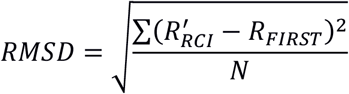

where *N* is the number of residues in the protein, 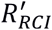 is the local rigidity computed with RCI and rescaled as described above, and *R_FIRST_* is the local rigidity computed with FIRST. The numerical values of correlation score and RMSD score are reported as the percentiles relative to a reference dataset formed of structures from the CNS and CNW datasets from the RECOORD recalculated structure database, which provide a representative selection of different fold types, before and after explicit solvent refinement.

### Dataset of comparable X-ray and NMR structures

To build a dataset of comparable X-ray and NMR structures we made use of the set of X-ray structures that were used to train the SHIFTX2 program for predicting chemical shifts^43^. This set comprises 197 high-resolution and high-quality structures which are representative of different fold types. We extracted structures which had corresponding NMR structures in the PDB, and backbone chemical shift completeness of at least 75%. Our final dataset consisted of 80 X-ray structures and 121 corresponding NMR structures for 68 different proteins. PDB and BMRB IDs are provided in supplementary information.

X-ray structures required some processing. If the structure contained multiple conformations (typical in high resolution X-ray structures), then we only considered the first of these as they appeared in the PDB file. Missing atoms and small breaks in the protein structure were identified using an in-house program and fixed using MODELLER^57^. MODELLER was also used to replace nonstandard residues related to conditions required for crystallisation (e.g. selenomethionine was replaced with methionine). Structures were protonated using REDUCE with the option to optimize adjustable groups^58^.

### Program speed and availability

We anticipate that the method will be of interest to a wide range of users who are interested in making use of NMR structures and would like some idea of their accuracy; and also to NMR spectroscopists who are doing a protein structure calculation and want to check their progress. The program and associated documentation can therefore be downloaded from github.com/nickjf/ANSURR. A typical calculation on an ensemble of 20 models for a 150-residue protein takes less than a minute.

## Acknowledgements

We thank the Biotechnology and Biological Science Research Council (BBSRC) for funding to N. J. F. (BB/P020038/1), and CREST, Japan Science and Technology Agency (JST) and PRISM for funding to A. S.

## Author contributions

M. P. W. and A. S. conceived the study. N. J. F. wrote the code and did the analysis. All authors wrote the manuscript.

