## Supplementary Information for "A method for validating the accuracy of NMR protein structures"

**Supplementary Figure 1.** Frequency distributions of output values for RCI (1a) and FIRST (1b). RCI values comprise residues from 7499 BMRB chemical shift entries related to NMR solution structures from the PDB. FIRST values comprise residues from 334 ensembles from the RECOORD CNW dataset. Figure 1c compares these frequency distributions after rescaling RCI as outlined in the methods section of the manuscript. Overall, there is reasonable agreement between the frequency distributions. As we use the CNW dataset for much of our analyses, we decided to compare the frequency distribution of FIRST for a random selection of 408 NMR solution structures from the PDB (1d). Here we also see reasonable agreement, although less good than for the CNW dataset. This is expected as structures in the CNW dataset are likely to be more accurate than a random structure from the PDB.

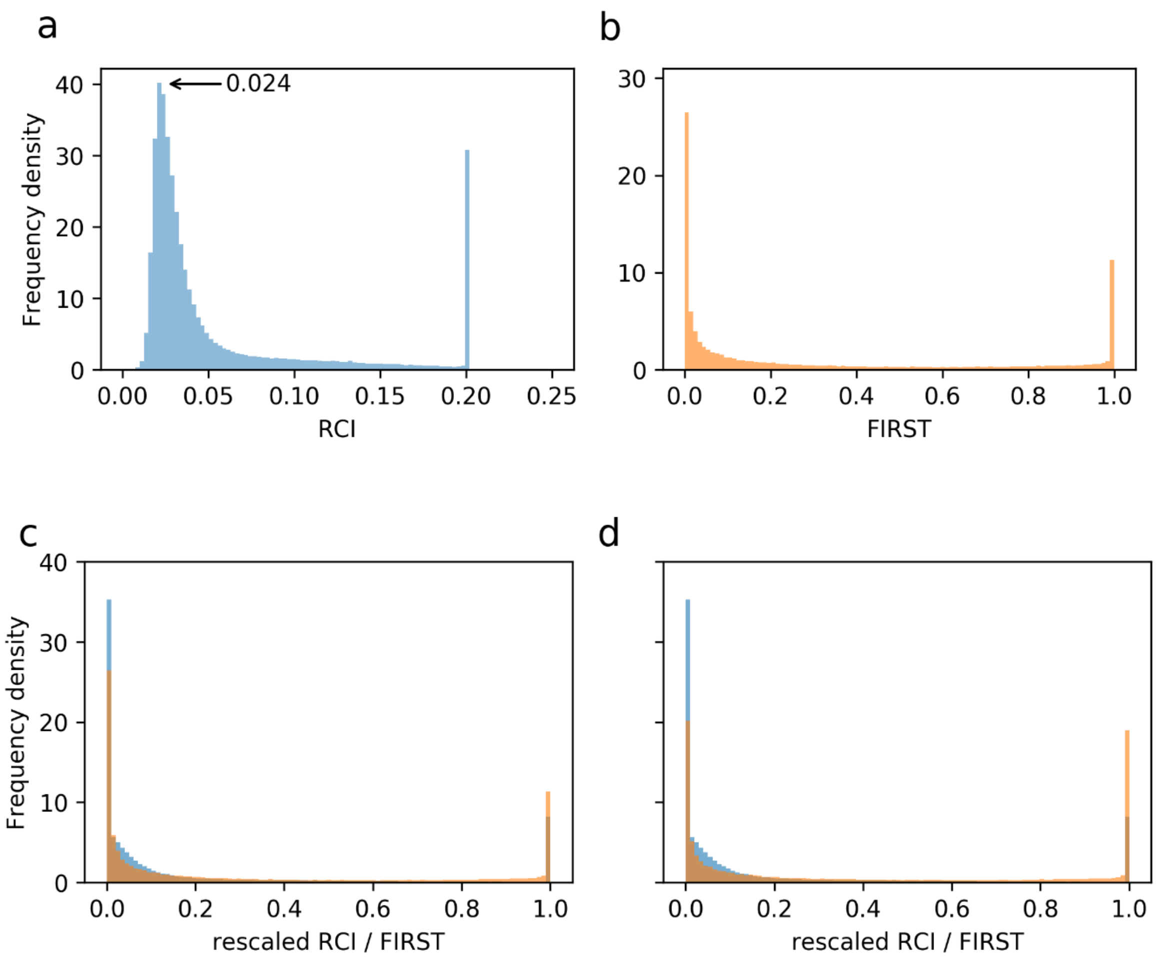

**Supplementary Table 1.** PDB IDs for 173 ensembles from the RECOORD CNS and CNW datasets with chemical shift completeness ≥ 75 %.

| | | | |
|---|---|---|---|
| 1A5J | 1F2H | 1JQR | 1N88 |
| 1B22 | 1F3Y | 1JR6 | 1N91 |
| 1B2T | 1F43 | 1JRM | 1ND9 |
| 1B4R | 1F53 | 1JW2 | 1NOR |
| 1B64 | 1FMM | 1JW3 | 1NR3 |
| 1B75 | 1FO7 | 1JWE | 1NWB |
| 1BCN | 1FR0 | 1JYT | 1NWV |
| 1BJX | 1G03 | 1JZU | 1NXI |
| 1BLR | 1G11 | 1K0S | 1NY4 |
| 1BNO | 1G4F | 1K0X | 1NY9 |
| 1BO0 | 1G6J | 1K1C | 1NZP |
| 1BQZ | 1G9L | 1K5O | 1O1W |
| 1C05 | 1GA3 | 1K8B | 1OCA |
| 1C06 | 1GH5 | 1K8O | 1OMT |
| 1C3T | 1H2O | 1K9C | 1OMU |
| 1C54 | 1H3Z | 1KFT | 1ONB |
| 1CEJ | 1H95 | 1KHM | 1OP1 |
| 1CEY | 1HS7 | 1KKG | 1PBU |
| 1CFC | 1HX7 | 1KMA | 1PFL |
| 1CFE | 1HY8 | 1KMD | 1PLO |
| 1CMZ | 1I42 | 1KOT | 1PN5 |
| 1COK | 1I5J | 1KRW | 1PUX |
| 1CX1 | 1ICH | 1KTM | 1Q27 |
| 1CZ5 | 1IEH | 1L1P | 1Q2N |
| 1D1N | 1IFW | 1L7B | 1Q59 |
| 1D3Z | 1IQO | 1L7Y | 1QHK |
| 1D8B | 1IRZ | 1LG4 | 1QND |
| 1DC7 | 1ITF | 1LM0 | 1QXF |
| 1DCJ | 1IX5 | 1LS4 | 1SPY |
| 1DD2 | 1J0T | 1M12 | 1SSN |
| 1DMO | 1J8I | 1M2F | 1SUH |
| 1DOQ | 1J8K | 1M5Z | 1TBD |
| 1DS9 | 1JAS | 1M7T | 1UD7 |
| 1DU2 | 1JDQ | 1M94 | 1XNA |
| 1DV5 | 1JE4 | 1MG8 | 2CPB |
| 1E17 | 1JE9 | 1MJD | 2CPS |
| 1E41 | 1JFJ | 1MKE | 2EZA |
| 1EGX | 1JFN | 1MP1 | 2EZH |
| 1EIH | 1JGK | 1MUT | 2MOB |
| 1EWW | 1JH3 | 1MZK | 3BDO |
| 1EZA | 1JI8 | 1N3G | 3PDZ |
| 1EZO | 1JJG | 1N4I | |
| 1EZP | 1JNS | 1N6U | |
| 1EZY | 1JOR | 1N6Z | |

**Supplementary Table 2.** 79 models from the RECOORD dataset used to generate decoys.

| PDB ID | Model number |
|---|---|
| 1A5J | 18 |
| 1B2T | 1 |
| 1B4R | 18 |
| 1B64 | 12 |
| 1B75 | 16 |
| 1BJX | 4 |
| 1BQZ | 11 |
| 1CFC | 20 |
| 1CMZ | 4 |
| 1COK | 8 |
| 1CX1 | 18 |
| 1DC7 | 4 |
| 1DD2 | 12 |
| 1DMO | 22 |
| 1DOQ | 3 |
| 1E17 | 22 |
| 1EGX | 21 |
| 1EWW | 13 |
| 1EZA | 11 |
| 1F2H | 8 |
| 1F43 | 21 |
| 1F53 | 14 |
| 1FO7 | 1 |
| 1G03 | 13 |
| 1G11 | 2 |
| 1G4F | 13 |
| 1GA3 | 7 |
| 1GH5 | 10 |
| 1H95 | 17 |
| 1HX7 | 15 |
| 1ICH | 17 |
| 1IEH | 20 |
| 1IFW | 14 |
| 1IRZ | 22 |
| 1ITF | 13 |
| 1IX5 | 1 |
| 1J8I | 10 |
| 1J8K | 5 |
| 1JE9 | 8 |
| 1JJG | 20 |

| PDB ID | Model number |
|---|---|
| 1JNS | 15 |
| 1JOR | 15 |
| 1JR6 | 9 |
| 1JWE | 11 |
| 1JZU | 5 |
| 1K0X | 24 |
| 1K5O | 1 |
| 1K9C | 4 |
| 1KKG | 7 |
| 1KMA | 9 |
| 1KMD | 22 |
| 1KRW | 22 |
| 1L7Y | 7 |
| 1LG4 | 14 |
| 1LM0 | 13 |
| 1M12 | 10 |
| 1M2F | 12 |
| 1M7T | 10 |
| 1MJD | 4 |
| 1MP1 | 16 |
| 1N3G | 2 |
| 1N4I | 14 |
| 1N6U | 23 |
| 1N6Z | 19 |
| 1NOR | 20 |
| 1NXI | 5 |
| 1PBU | 10 |
| 1PFL | 21 |
| 1PN5 | 2 |
| 1Q2N | 7 |
| 1SPY | 3 |
| 1SUH | 10 |
| 1TBD | 3 |
| 1UD7 | 4 |
| 1XNA | 5 |
| 2CPB | 9 |
| 2EZA | 3 |
| 2EZH | 4 |
| 3PDZ | 19 |

**Supplementary Figure 2.** Validation scores computed for each target structure in supplementary table 2 (black asterisk) and 300 decoy structures (circles colored according to GDT). Each plot shows one protein, indicated by PDB ID and the percentage of α-helix and β-sheet in the target structure.

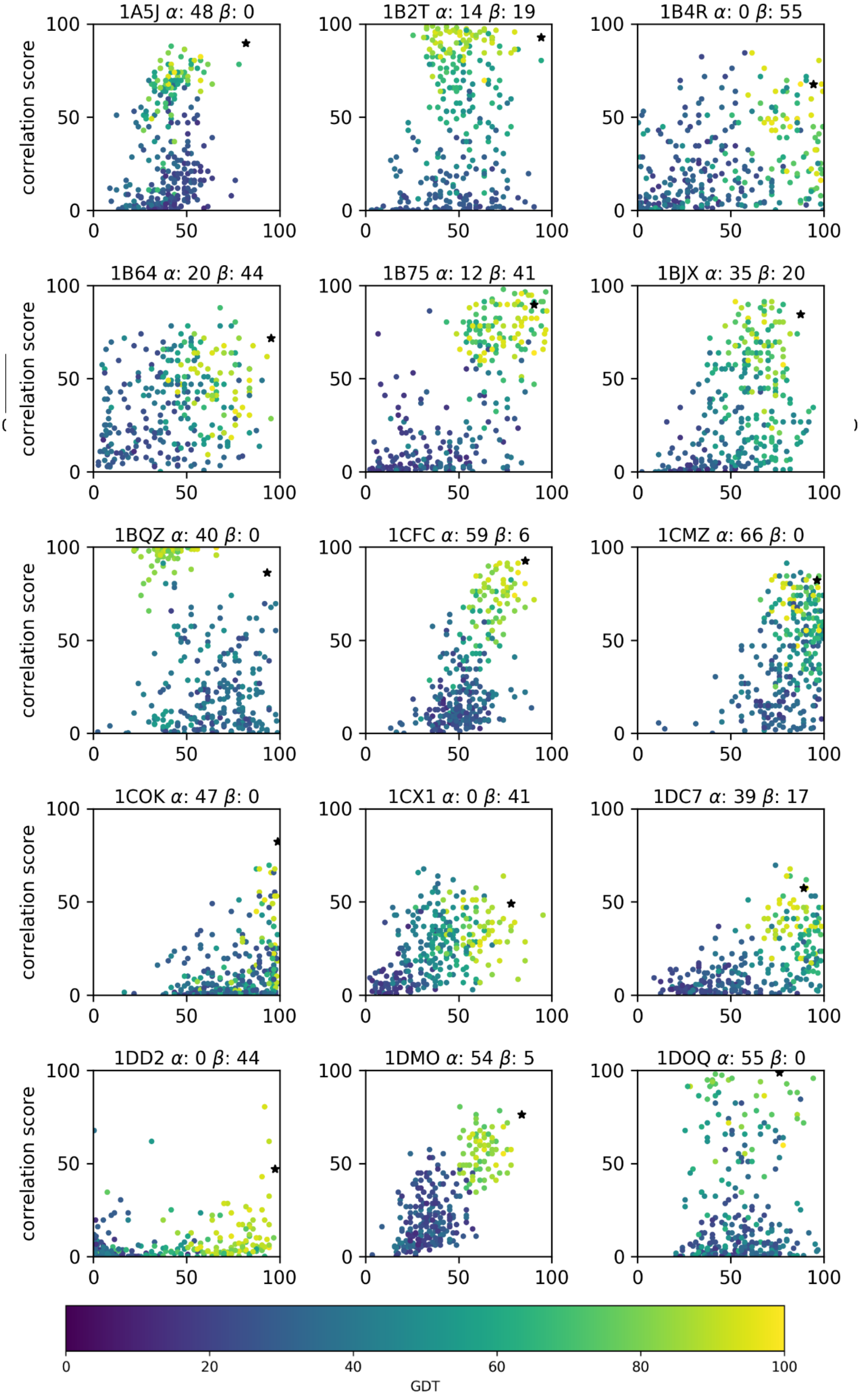

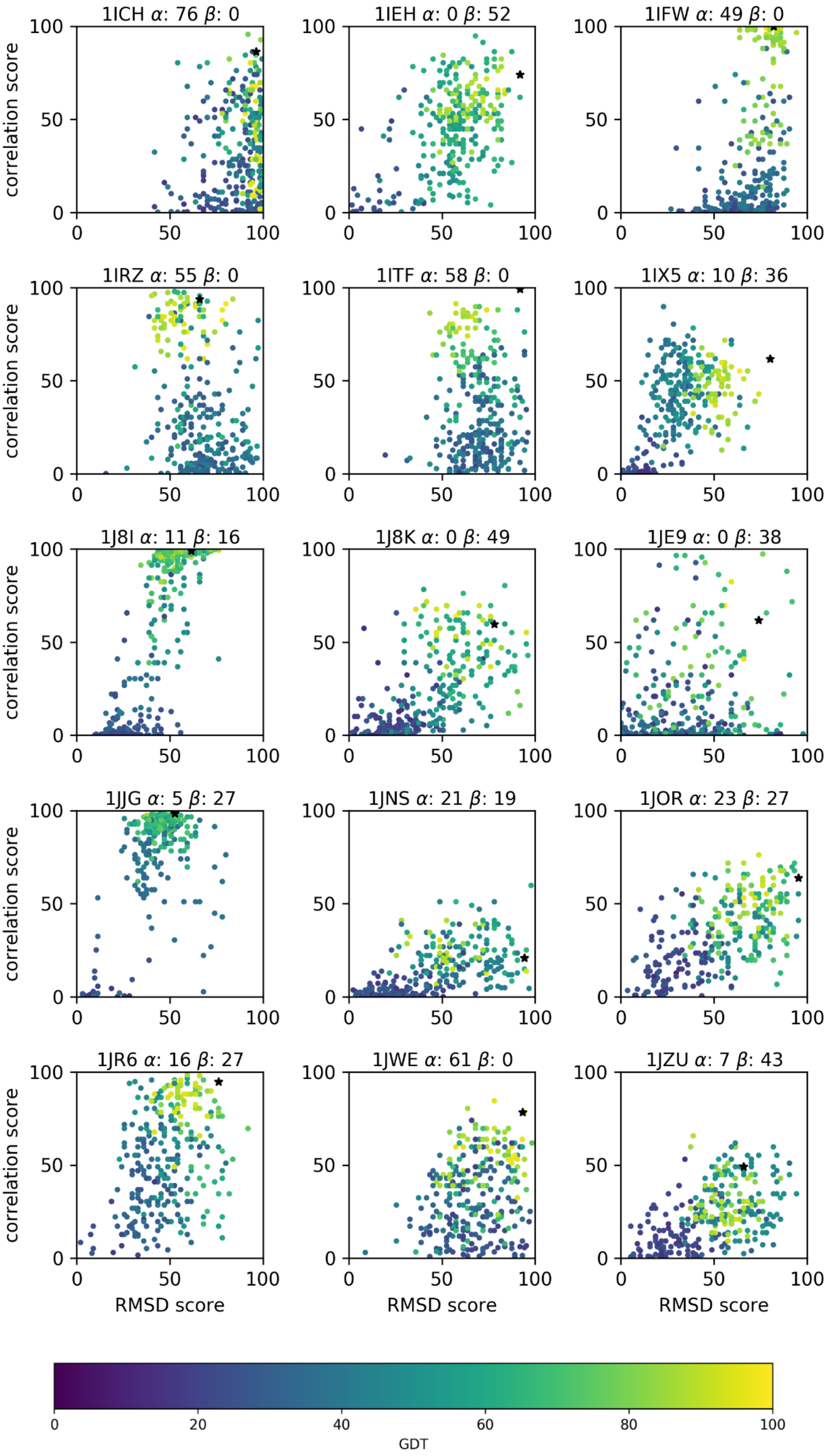
1ICH α: 76 β: 0
1IEH α: 0 β: 52
1IFW α: 49 β: 0
1IRZ α: 55 β: 0
1ITF α: 58 β: 0
1IX5 α: 10 β: 36
1J8I α: 11 β: 16
1J8K α: 0 β: 49
1JE9 α: 0 β: 38
1JJG α: 5 β: 27
1JNS α: 21 β: 19
1JOR α: 23 β: 27
1JR6 α: 16 β: 27
1JWE α: 61 β: 0
1JZU α: 7 β: 43
correlation score
RMSD score
0
50
100
0
20
40
60
80
100
GDT

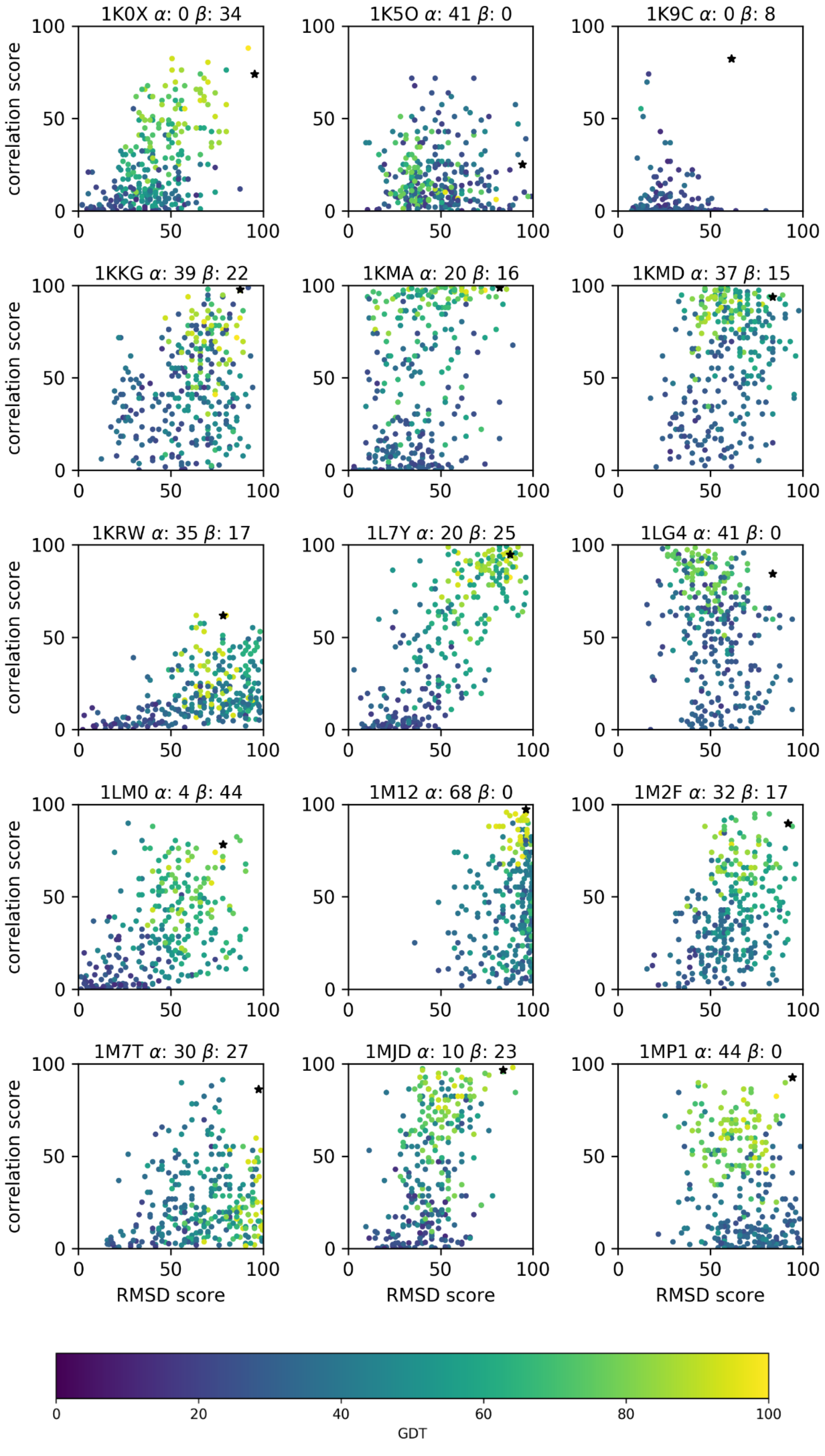

1K0X α: 0 β: 34
1K5O α: 41 β: 0
1K9C α: 0 β: 8
1KKG α: 39 β: 22
1KMA α: 20 β: 16
1KMD α: 37 β: 15
1KRW α: 35 β: 17
1L7Y α: 20 β: 25
1LG4 α: 41 β: 0
1LM0 α: 4 β: 44
1M12 α: 68 β: 0
1M2F α: 32 β: 17
1M7T α: 30 β: 27
1MJD α: 10 β: 23
1MP1 α: 44 β: 0
correlation score
RMSD score
GDT

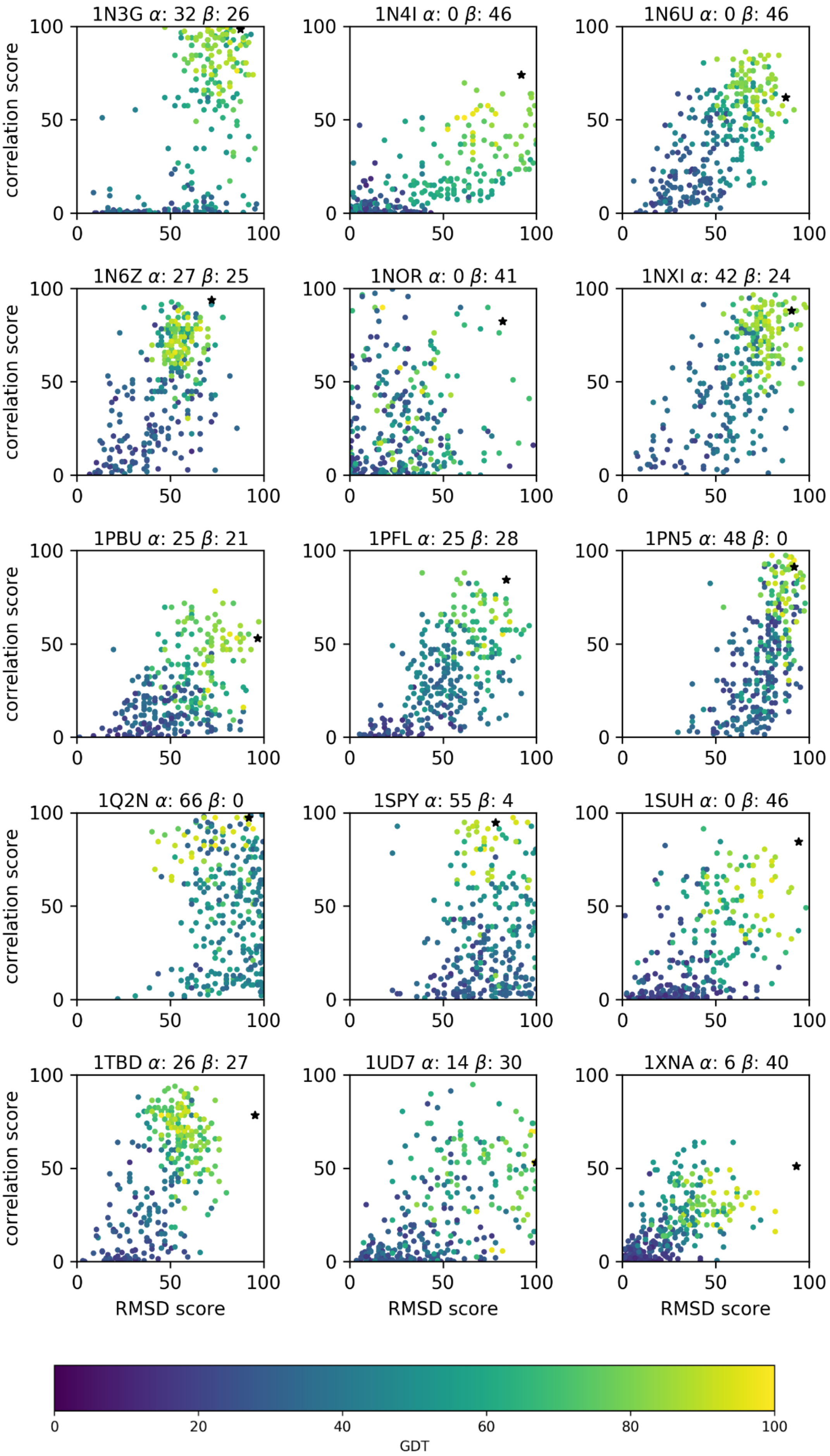

1N3G α: 32 β: 26
1N4I α: 0 β: 46
1N6U α: 0 β: 46
1N6Z α: 27 β: 25
1NOR α: 0 β: 41
1NXI α: 42 β: 24
1PBU α: 25 β: 21
1PFL α: 25 β: 28
1PN5 α: 48 β: 0
1Q2N α: 66 β: 0
1SPY α: 55 β: 4
1SUH α: 0 β: 46
1TBD α: 26 β: 27
1UD7 α: 14 β: 30
1XNA α: 6 β: 40
correlation score
RMSD score
0
50
100
GDT
0
20
40
60
80
100

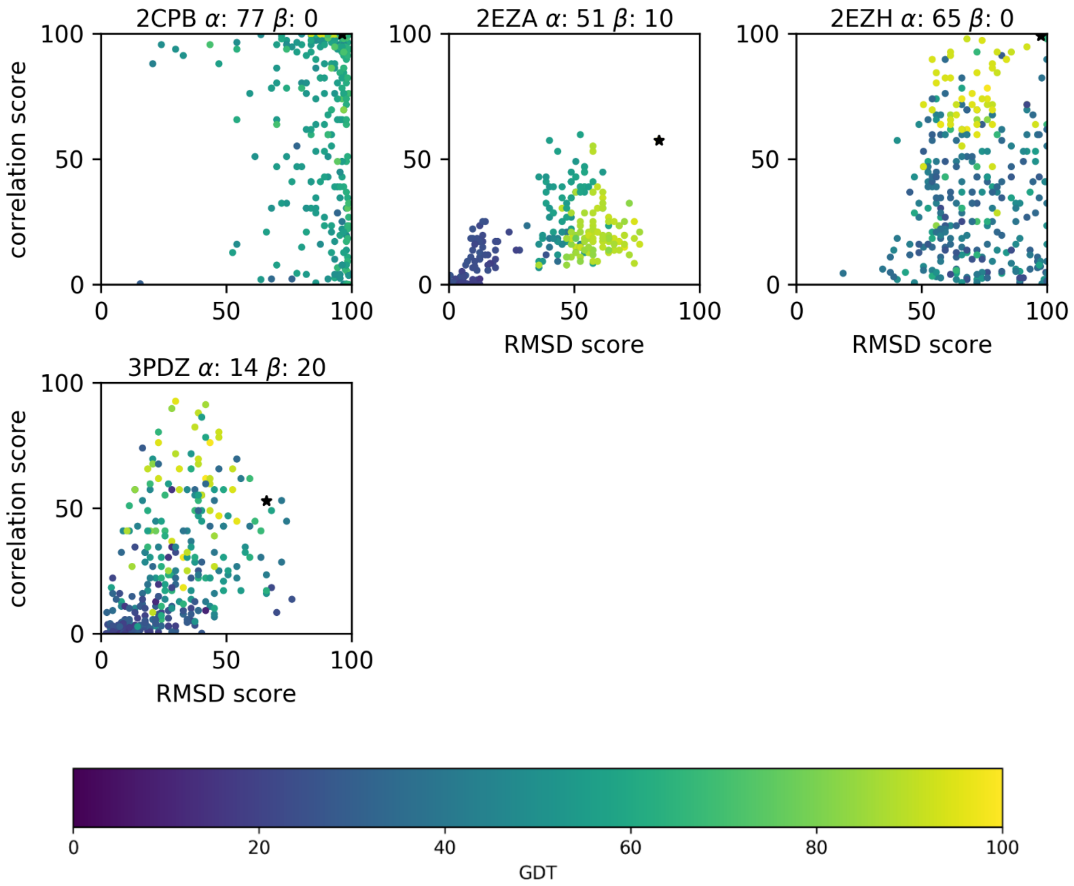

2CPB $\alpha$: 77 $\beta$: 0
2EZA $\alpha$: 51 $\beta$: 10
2EZH $\alpha$: 65 $\beta$: 0
3PDZ $\alpha$: 14 $\beta$: 20
correlation score
RMSD score
0
50
100
0
20
40
60
80
100
GDT

**Supplementary Table 3.** BMRB and PDB IDs for a set of comparable X-ray and NMR structures.

| BMRB ID | X-ray PDB ID | NMR PDB ID |
|---|---|---|
| 4019 | 1PLC | 1TKW |
| 4031 | 1RUV,3RN3 | 2AAS |
| 4039 | 1RSY | 1BYN,2K45,2K4A,2K8M |
| 4052 | 1SNC | 1JOK,1JOO,1JOQ,1JOR |
| 4064 | 1HFC | 1AYK,2AYK,3AYK,4AYK |
| 4082 | 1FIL | 1PFL |
| 4094 | 2B8X | 1BBN,1BCN,1ITI |
| 4115 | 1EMV | 1E0H,1IMP,1IMQ,2K5X |
| 4162 | 1EPF | 2NCM |
| 4186 | 1CBS | 1BLR |
| 4198 | 1EZ3 | 1BR0 |
| 4202 | 1HL5 | 1BA9,1KMG,1RK7 |
| 4259 | 1RGE,1UCK | 1C54 |
| 4296 | 1MJC | 2L15 |
| 4317 | 1AIL | 1NS1 |
| 4340 | 1JV4,1YP7 | 1DF3 |
| 4342 | 1EKG | 1LY7 |
| 4354 | 1LAX,1FQA | 1EZO,1EZP,2H25,2KLF |
| 4371 | 1ONC | 1PU3 |
| 4378 | 1G8I | 2LCP |
| 4401 | 1TOP,1NCX | 1BLQ,1SKT,1TNP,1TNQ,<br>1ZAC |
| 4421 | 1GXQ | 1QQI |
| 4425 | 1BDO | 3BDO |
| 4438 | 1C44 | 1QND |
| 4472 | 1U8T | 1DJM |
| 4553 | 1XUO | 1DGQ |
| 4566 | 1AYF | 1L6U,1L6V |
| 4717 | 1F46,1Y2G | 1F7W,1F7X |
| 4797 | 1IAZ | 1KD6 |
| 4840 | 2CDN | 1P4S |
| 4857 | 2A0B | 1FR0 |
| 4964 | 1A2P,1BRI | 1BNR,1FW7 |
| 5058 | 1GNU | 1KOT |
| 5081 | 1B2V | 1YBJ |
| 5142 | 1IWT,1LZ1 | 1IY3,1IY4 |
| 5194 | 1TJM | 1K5W |
| 5206 | 1MHO | 1UWO |
| 5211 | 1D4T | 1KA7 |
| 5220 | 1I1J | 1K0X |
| 5275 | 1KQR | 1KRI |
| 5299 | 2VFX | 1KLQ |
| 5387 | 1UBQ | 1D3Z,1G6J |
| 5471 | 1P7T | 1Y8B |
| 5485 | 1W41,1H7M | 1GO0 |
| 5712 | 1IPB | 2GPQ |
| 5756 | 1N0S | 1T0V |
| 5792 | 1EW4 | 1SOY |

| BMRB ID | X-ray PDB ID | NMR PDB ID |
|---|---|---|
| 5843 | 1Q4R | 1Q53 |
| 5898 | 1UOH | 1TR4 |
| 5921 | 1VC1 | 1SBO |
| 6075 | 1JL3 | 1Z2D,1Z2E |
| 6122 | 1SMX,1SN8 | 1SLJ |
| 6231 | 1UV0 | 2GO0 |
| 6375 | 1U07 | 1XX3 |
| 6503 | 1F2F | 2JYQ |
| 6504 | 1CNR | 1CCM,1CCN,1YV8,1YVA, 2EYA,2EYB,2EYC,2EYD |
| 6541 | 1CLL,1MXE,2F3Y | 1CFC,1CFD,1CFF,1CKK, 1MUX,1NWD,1SY9,1X02, 2JZI,2K0E,2K0F,2KDU, 2KNE,2L53 |
| 6699 | 4ICB | 1N65,2MAZ |
| 6754 | 1QAV | 1Z86 |
| 6776 | 1UJ8 | 2BZT |
| 4070 | 1ZE3 | 1BF8 |
| 6090 | 1FF3 | 2GT3 |
| 6876 | 2NNR | 2FO8 |
| 6922 | 2D3D | 2FE9 |
| 6923 | 2AWG | 2MF9 |
| 6932 | 1KBL | 2FM4 |
| 6980 | 2D58 | 2G2B |
| 15084 | 1TW4 | 1MVG,1ZRY,2JN3,2K62 |

**Validation of lysozyme X-ray structures obtained at 100 K and 278 K.**

Here we compare X-ray structures of hen egg-white lysozyme obtained at 100 K and 278 K. We chose a set of structures deposited by the same research group for which crystal preparation and data collection were performed by following the same protocol. Structures were downloaded from the PDB (PDB IDs for structures obtained at 100 K: 5KXK, 5KXL, 5KXM, 5KXN; and at 278 K: 5KXO, 5KXP, 5KXR, 5KXS ,5KXT, 5KXW, 5KXX, 5KXY, 5KXZ, 5KY1) and processed in the same way as described in Methods - Dataset of comparable X-ray and NMR structures. Backbone chemical shifts were extracted from two sets deposited to the BMRB (H shifts from BMRB ID 4565 and N/C shifts from BMRB ID 4851). We then validated each structure using ANSURR. The results presented in supplementary Fig. 3.1 show that the structures obtained at 278 K are better. That is to say, that flexibilities predicted for those structures are a better match to those predicted from chemical shifts obtained in solution at room temperature. These results suggest that X-ray structures obtained at cryogenic temperatures are more rigid that at room temperature. In figures 3.2 and 3.3, we compare flexibility predicted by RCI and FIRST for each structure at 100 K and 278 K, respectively. Noticeable is that for structures obtained at 100 K there are many missing peaks in flexibility that should be present according to RCI. Structures obtained at 278 K have many more of these peaks in flexibility. Backbone superposition of the structures shows that the fold is essentially identical at both temperatures (Fig. 3.4). From inspection of the structures, the over-rigidification of structures obtained at 100 K manifests as additional and/or slightly stronger hydrogen bonds in loop regions.

**Supplementary Fig. 3.1.** Validation scores for lysozyme X-ray structures obtained at 100 K (blue) and 278 K (orange).

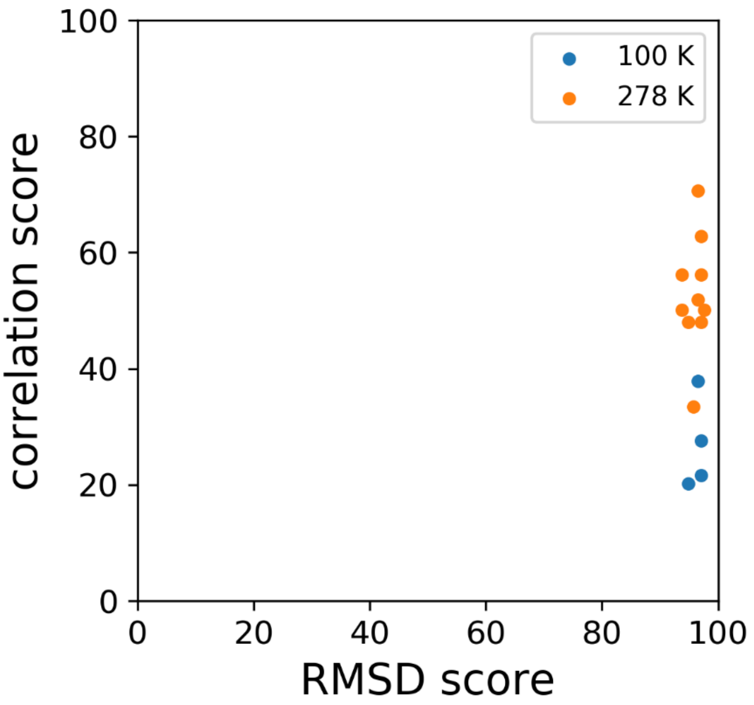

**Supplementary Fig. 3.2.** Comparison of flexibility predicted by RCI and FIRST for lysozyme X-ray structures obtained at 100 K.

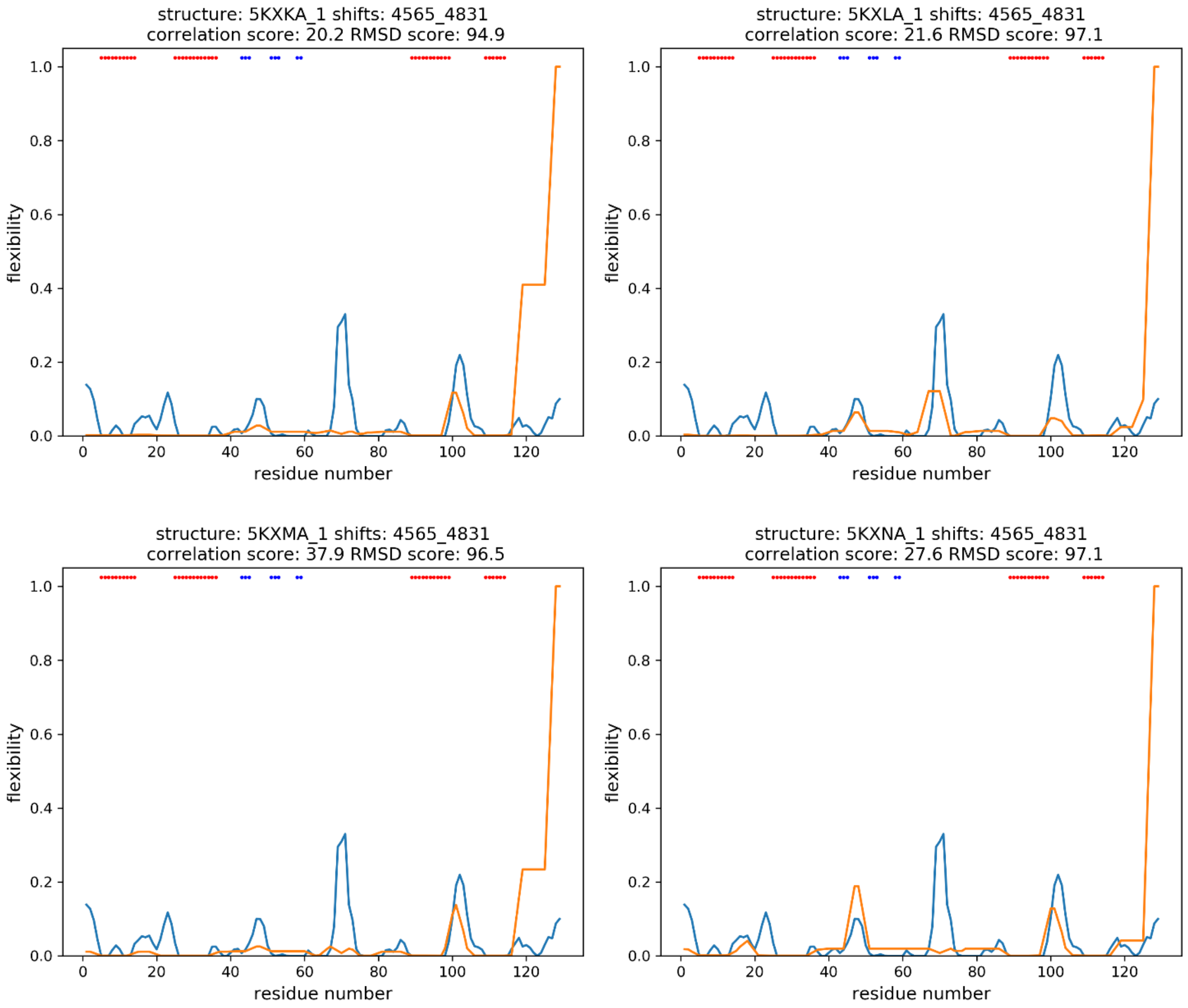

**Supplementary Fig. 3.3.** Comparison of flexibility predicted by RCI and FIRST for lysozyme X-ray structures obtained at 278 K. Continued on the next page.

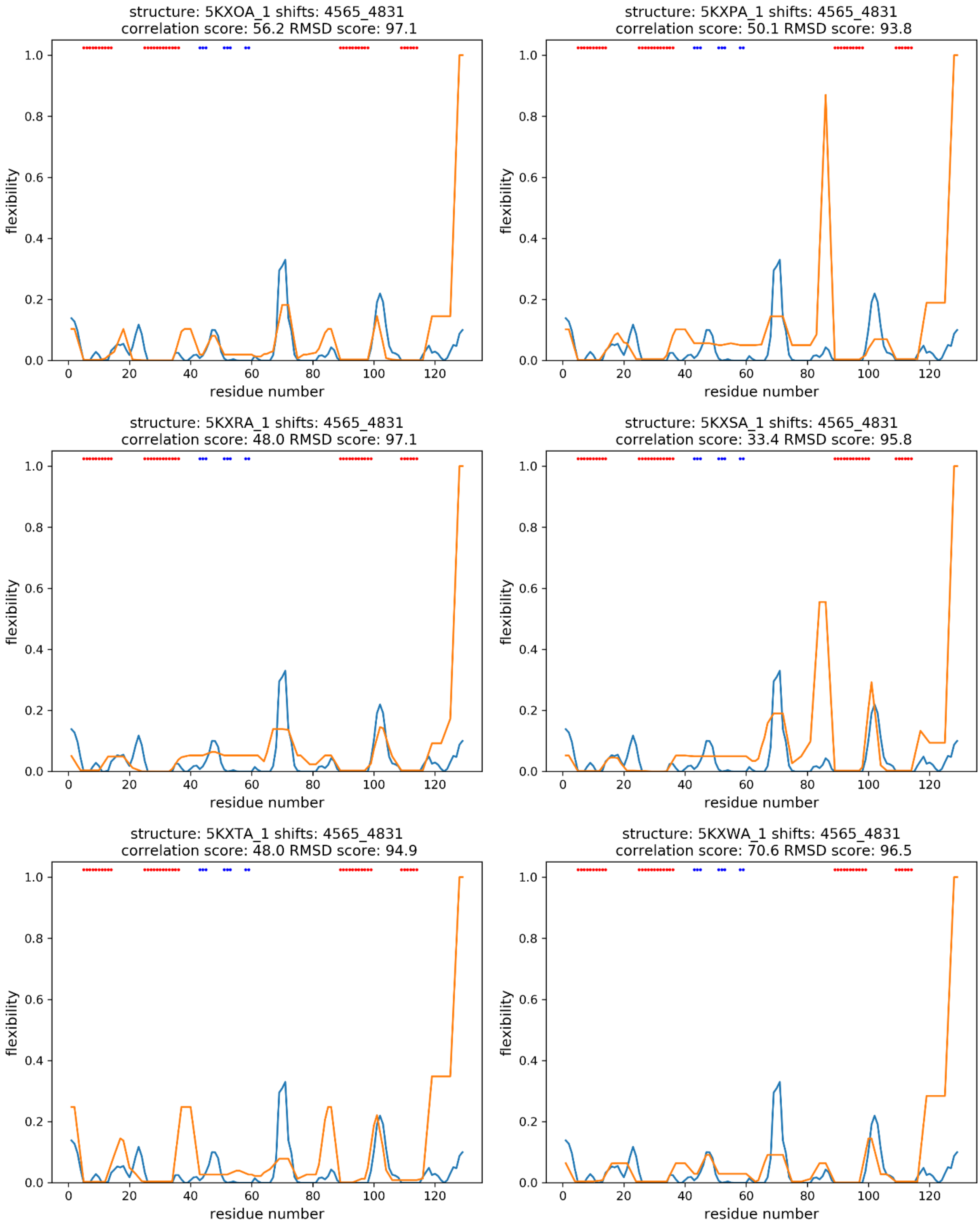

**Supplementary Fig. 3.3 continued**. Comparison of flexibility predicted by RCI and FIRST for lysozyme X-ray structures obtained at 278 K.

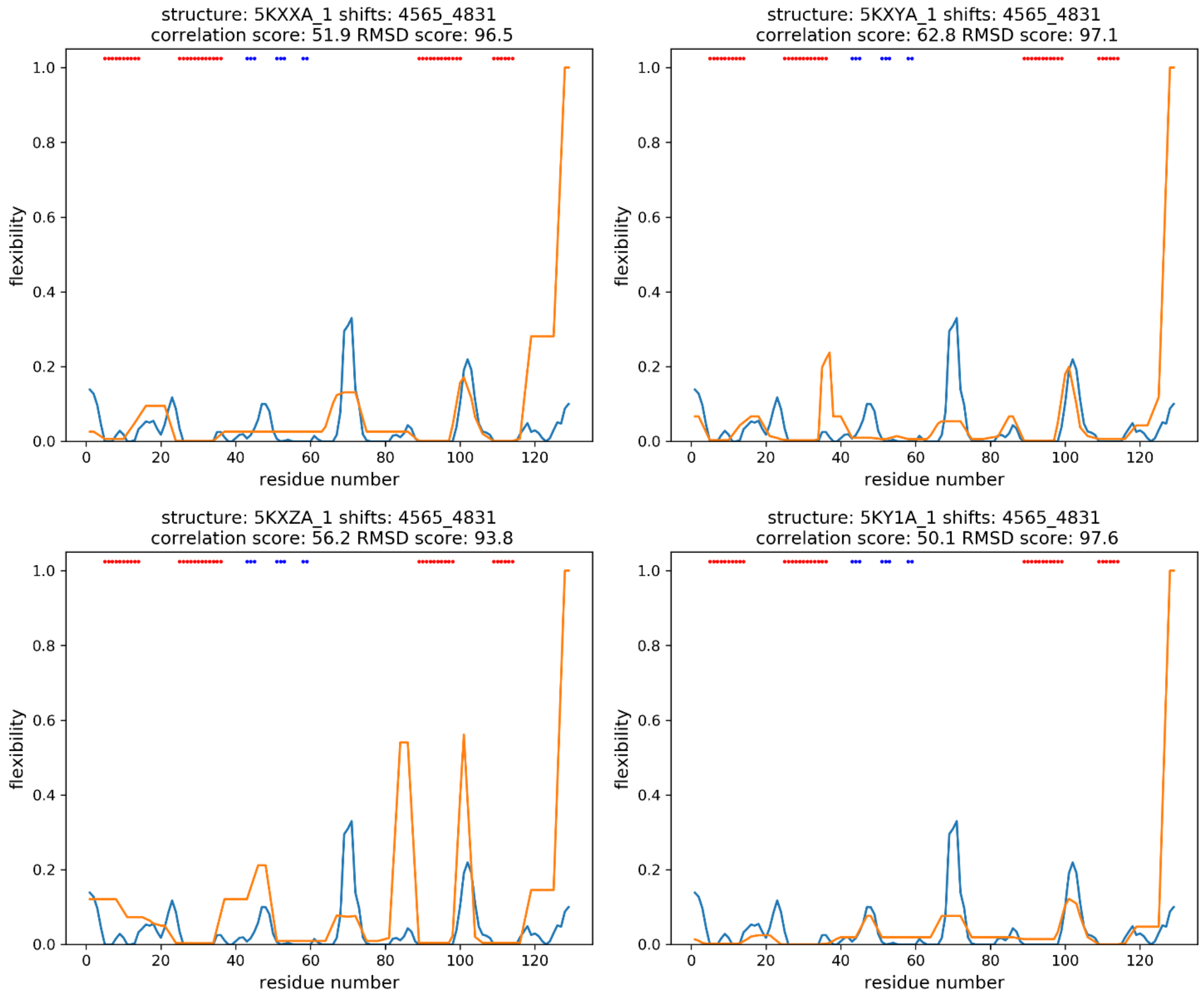

**Supplementary Fig. 3.4.** Backbone superposition of lysozyme X-ray structures obtained at 100 K (blue) and 278 K (orange).

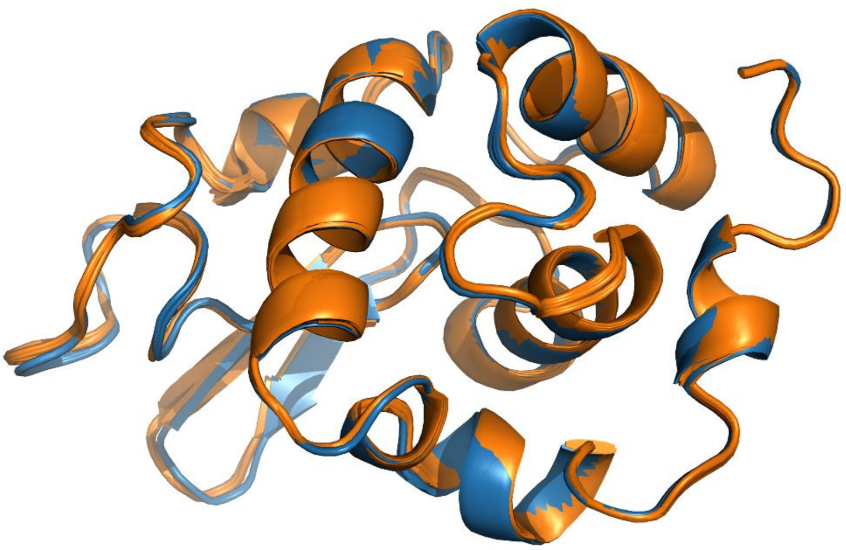
